# The Value of Multi-Year Sampling for Detecting Fine-Scale Population Genetic Structure in Marine Fishes: A Case Study of Juvenile Southern Flounder

**DOI:** 10.64898/2026.04.24.720543

**Authors:** Sydney P. Harned, Jamie L. Mankiewicz, Russell J. Borski, John Godwin, Martha O. Burford Reiskind

**Author notes:** Corresponding author: Sydney Harned.

## Abstract

Understanding population structure is critical for effective fisheries management in species with complex life histories and variable recruitment. Southern flounder (*Paralichthys lethostigma*) is a valuable flatfish species with declining populations in the Southeast United States. Improved management may depend on a better understanding of fine-scale and temporal population genetic structure in this region; however, such structure remains poorly characterized. To address our lack of understanding of the spatial and temporal population structure of this important species, we used double digest reduced-representation genome sequencing (ddRADSeq) on juveniles from estuaries in North Carolina and Texas between 2014 and 2023. We found significant genetic differentiation between the Gulf of Mexico and Atlantic populations, supporting the management of these regions as distinct stocks. By contrast, we detected significant variance in genetic structure within Texas and North Carolina populations that was not consistent across sampling years between estuaries in close proximity. The population genetic structure of southern flounder suggests significant, temporally variable genetic differences within estuarine locations that may result from variation in larval dispersal and recruitment patterns. Our findings highlight the value of integrating fine-scale, multi-year genetic data to capture temporal dynamics and avoid misleading conclusions based on single-year or broad-scale sampling.

## 1. Introduction

The assessment of genetic structure and population connectivity is fundamental for understanding recruitment dynamics in marine species (Cowen et al., 2007; Tova Verba et al., 2023). However, high dispersal potential and pelagic larval stages can make genetic population structure difficult to resolve (Palumbi 2003; Selkoe et al. 2008). While high dispersal often leads to high connectivity in marine systems, many marine species exhibit population structure despite this potential (Parrondo et al. 2022). Advances in genomic sequencing now enable the analysis of tens of thousands of genomic markers, improving the detection of fine-scale population structure found in these dynamic marine systems (Parrondo et al. 2022). These advances have improved our understanding of population structure across space, but to fully characterize population structure in a region requires replication in time as well. While pooling samples across years can provide a useful, time-averaged view of connectivity, this approach may obscure temporal variability driven by stochastic larval dispersal and variable retention. Genetic patterns can be patchy and inconsistent across cohorts and may not be captured in a single sampling year or when samples are combined (Christie et al. 2010; Hedgecock et al. 2007). By focusing on juveniles, which represent discrete recruitment events, temporal sampling allows for more precise inference of cohort-specific structure and connectivity in marine species characterized by high dispersal potential.

Temporal population genomic studies can provide insight into recruitment and spawning stock dynamics in marine fishes. These dynamics include temporal fluctuations in reproductive success among spawning groups, including sweepstakes reproductive events, variable larval survival, and shifting contributions from different spawning locations or cohorts (Hedgecock & Pudovkin 2011; Christie et al. 2010). From a management perspective, this information is essential for understanding how the breeding population contributes to recruitment, evaluating the stability of spawning stock composition, and assessing whether local populations function as discrete units or are replenished by variable sources over time. Consequently, incorporating temporal genomic data strengthens stock assessment and informs management strategies by linking genetic patterns directly to recruitment variability and spawning stock dynamics.

Southern flounder (*Paralichthys lethostigma*) is a widely distributed coastal flatfish occurring throughout the U.S. Southeast Atlantic and Gulf of Mexico (Flowers et al. 2019). The species exhibits life history traits common to marine fishes, including offshore spawning and a pelagic larval stage, which promote high dispersal and connectivity across its range (Fig. 1 inset; Chrisp et al. 2023). Previous genetic studies using allozymes/microsatellites and mtDNA/AFLPs supported strong Atlantic–Gulf differentiation and weak or inconsistent structure within the U.S. Atlantic, with inferences largely drawn from broad spatial sampling of only a few sites at one point of time (Blandon et al. 2001; Anderson & Karel 2012). A genetic analysis using mitochondrial DNA and AFLPs further identified weak structure and evidence of basin-wide mixing within the U.S. Atlantic, without explicit evaluation of temporal stability (Wang et al. 2015). However, a recent study applying next-generation sequencing approaches found some evidence of estuarine-scale structure in southern flounder in the Gulf and Atlantic using samples pooled across years and found that genetic heterogeneity may be driven by environmental variation rather than geographic distance (O’Leary et al. 2021). However, whether fine-scale genetic structure is present and consistently detectable across estuaries through time remains unresolved in both the Gulf and Atlantic populations. Building on this work, resolving fine-scale and potentially transient genetic structure within estuaries requires higher-resolution genomic data with temporal replication.

**Fig 1.**
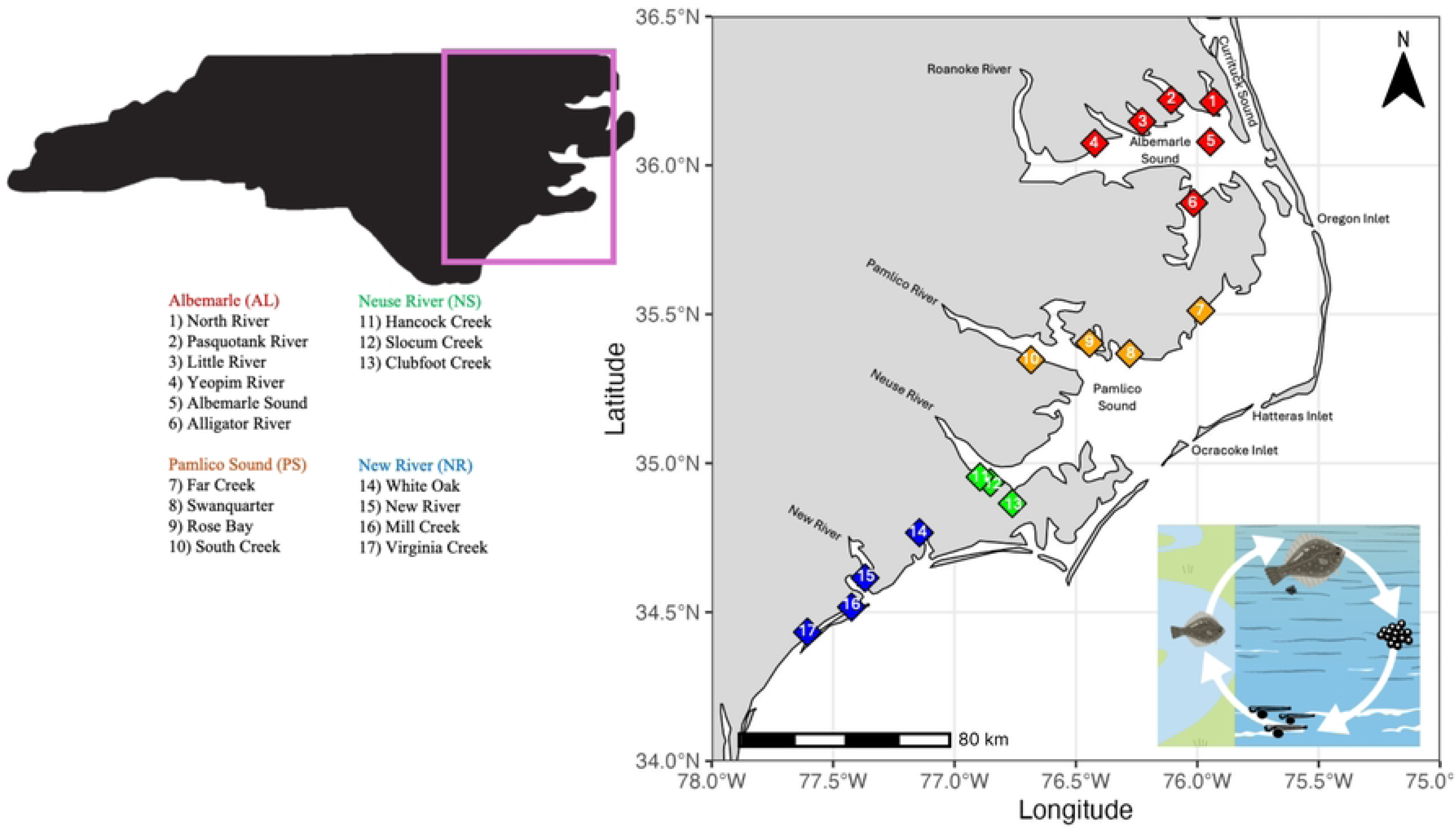
Map of sampling locations for southern flounder (*Paralichthys lethostigma*) in North Carolina estuaries. The inset highlights the study region within North Carolina. Sampling sites are grouped into four regions: Albemarle (AL; sites 1–6), Pamlico Sound (PS; sites 7–10), Neuse River (NS; sites 11–13), and New River (NR; sites 14–17). Numbered markers correspond to individual sampling locations listed in the legend and are color-coded by region. The figure in the right corner of the map depicts the southern flounder life cycle: juveniles spend 1-2 years in shallow estuarine environments before moving offshore for spawning. Eggs hatch after 2-3 days, and planktonic larvae spend 30-60 days in the water column before juveniles settle in estuaries.

To address this, we assess fine-scale and temporal patterns of population genetic structure in juvenile southern flounder in North Carolina and Texas estuaries using high-resolution genomic data obtained through ddRAD sequencing, providing insight into genetic patterns associated with recent recruitment. We chose to focus primarily on North Carolina, a region with relatively few fine-scale population genetic studies and a spatially complex intracoastal structure. Specifically, we examine whether genetic differentiation is detectable within and among estuaries and whether these patterns are consistent across multiple sampling years. Given the species’ high dispersal potential, we expect that fine-scale genetic structure would be weak and temporally variable. By evaluating fine-scale genetic structure across space and time, this study informs key assumptions about connectivity in highly dispersive marine species within heterogeneous estuarine systems and supports the delineation of biologically relevant management units.

## 2. Materials and Methods

### 2.1 Sample Locations

Samples for this study were collected from two Ocean Basins: The Southeast U.S. Atlantic (North Carolina; Fig. 1) and the Gulf of Mexico (Texas; Fig. 2). We sampled juvenile fish from estuarine environments where larval flounder settle and remain for 1-2 years before moving offshore for spawning (Flowers et al. 2019). Estuarine sampling locations within North Carolina were grouped into three regions— Pamlico Sound, Neuse River, and South of the New River — based on geographic barriers and environmental differences.

**Fig 2.**
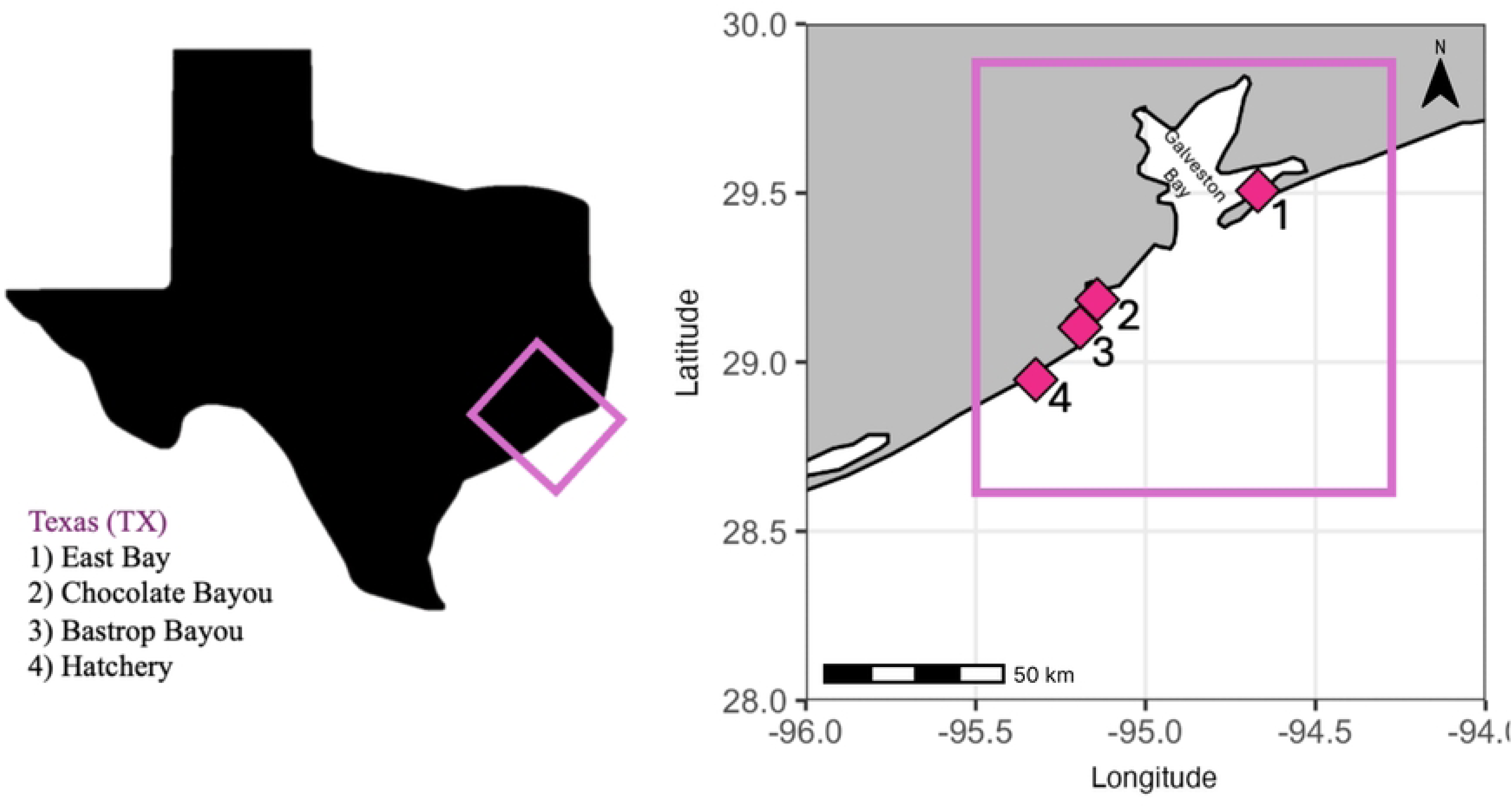
Map of sampling locations for southern flounder (*Paralichthys lethostigma*) in Texas. The inset highlights the study region along the upper Texas coast near Galveston Bay. Sampling sites include (1) East Bay, (2) Chocolate Bayou, (3) Bastrop Bayou, and (4) a hatchery population. Numbered markers correspond to the sampling locations shown on the map.

Each of the three regions in North Carolina are affected by different environmental and spatial factors (Fig 1). Pamlico Sound is an enclosed estuary system with the barrier islands to the East. Pamlico Sound has the lowest and most stable salinity and temperature of the regions due to limited tidal exchange and large freshwater inputs (Giese, Wilder and Parker 1985). While the Neuse River, our intermediate region, is connected to the ocean via Pamlico Sound, it has a higher rate of flushing than Pamlico and is more variable in temperature and salinity (Giese, Wilder and Parker 1985, Honeycutt et al. 2019). Our southernmost region, areas south of the New River (herein, New River), experiences higher average salinity and temperature, with greater fluctuations driven by tidal input and less freshwater flow (Giese, Wilder and Parker 1985; Honeycutt 2018; Honeycutt et al. 2019). In 2023, our expanded sampling included Albemarle Sound, a northern region with environmental conditions more closely aligned with Pamlico Sound; however, Albemarle Sound differs from Pamlico Sound as it is characterized by lower average temperatures, higher variation in freshwater inflow, and a single narrow inlet (the Oregon Inlet) connecting the Sound to the Atlantic Ocean (Copeland 1983). As sampling occurred in different locations from year-to-year, the sampling locations within each region vary. Specific sampling schema from each year can be found in supplemental Table S1.

### 2.2 Sample Collection

We obtained juvenile southern flounder from North Carolina Division of Marine Fisheries (NCDMF) biologists during the juvenile monitoring Program 120 (P120) estuarine trawl survey in 2014-2016, 2022, and 2023. The survey samples shallow, upper estuary locations using an otter trawl with a 3.2 m headrope, 6.4 mm bar mesh wings and body, and a 3.2 mm bar mesh cod end. The trawl is towed for 1 min at 1.1 m/s at each station. Sampling locations within four geographic regions are shown in Figure 1. In 2016, we supplemented NCDMF sample collection with our own sampling using an otter trawl or 2-meter-wide beam trawl depending on habitat (Honeycutt et al. 2019).

To assess broad genetic structure between ocean basins, we included juvenile samples collected in Texas in our analysis. We obtained Texas samples from Texas Parks and Wildlife in 2014, 2015 and 2016 from three locations: East Bay, Bastrop Bayou and Chocolate Bayo (Fig. 2). Fish were sampled using 1- and 2-meter-wide beam trawls pulled for 1-2 minutes at 1-2 knots. In areas where boat sampling was restricted, fish were sampled using trawls pulled by-hand for 20 meters. We also included samples from the Sea Center Texas Marine Fish Hatchery (Lake Jackson, TX) in our 2016 data set to assess the genetic similarity between hatchery and wild-caught Texas individuals.

### 2.3 Extraction, Sequencing and Genomic Library Building

We extracted genomic DNA from fin clips using Qiagen DNA Blood and Tissue kits, following the manufacturer’s instructions (Qiagen, Inc., Valencia, CA). We then quantified total genomic DNA with a Qubit fluorometer 4.0 (Invitrogen, Carlsbad, CA). We prepared ddRAD sequencing libraries with enzyme pair *SphI* and *MluCI* following the protocol detailed by Burford Reiskind et al. (2016). We digested genomic DNA with both enzymes in CutSmart buffer at 37°C for 3 hours. We then ligated adapters using T4 ligase, and incubated ligation reactions at 22°C for 2 hours followed by heat inactivation at 65°C for 20 minutes. We constructed libraries by pooling 200 ng of DNA from each sample into libraries containing 48 individuals. We then purified the pooled libraries using the Qiaquick PCR cleanup kit (Qiagen, Inc., Valencia, CA) and size selected for fragments between 360-475 bp using a BluePippin system (Sage Science, Beverly, MA). Following size selection, libraries were amplified and sequenced as single-end 90 bp fragments on the Illumina NovaSeq at the North Carolina State University Genomic Sequencing Laboratory (Raleigh, NC, USA).

### 2.4 Bioinformatic Pipeline

We used the program FastQC (Babraham Bioinformatics; http://www.bioinformatics.babraham.ac.uk/projects/fastqc/) to check the quality of reads, using a high phred score criterion (phred >33). We then processed barcodes as outlined in Burford Reiskind et al. (2016), using the *process_radtags* pipeline to filter and de-multiplex variable length barcodes in STACKS v. 1.24 (Catchen et al. 2011). We trimmed reads to identical lengths (90 bp) and used the *de novo* pipeline (*denovo.pl*) in STACKS to detect single nucleotide polymorphisms (SNPs) with the following parameters: *m* = 3 (minimum stack depth), *M* = 2 (mismatches allowed between loci within an individual), and *n* = 2 (mismatches allowed between loci combined in a catalog; Catchen et al. 2011). From the STACKS *de novo* pipeline, we identified 3,643,301 loci across all North Carolina and Texas samples with an average per-sample coverage of 22.2x. A reference genome was not available for this study, but *de novo* approaches reliably capture patterns of genetic differentiation in non-model organisms (Catchen et al. 2011).

Because a congener summer flounder (*P. dentatus*) overlaps in its range with southern flounder, and the juveniles are morphologically similar, we confirmed that all sampled individuals were *P. lethostigma* using Treemix v. 1.13 (Pickrell et al. 2012) to produce maximum-joining trees for each sampling year. To confirm branching patterns, we conducted a bootstrap analysis using 1000 replicates. We identified four individuals in 2023 that belonged to *P. dentatus.* These individuals were removed from the dataset.

### 2.5 Dataset Filtering and Transformation

We ran STACKS *populations* pipeline retaining loci present in at least one (-p 1) or two (-p 2) populations and requiring a minimum within-group genotyping rate of 60% (-r 0.6). We retained one SNP per rad locus (--write-random-snp) and excluded loci with high heterozygosity (--max-observed-het 0.8).

We then filtered in PLINK (PLINK v.1.19; http://pngu.mgh.harvard.edu/purcell/plink/) for minor allele frequency (maf 0.01) and locus missingness (geno 0.5). Because different analyses require different assumptions of locus sharing and data completeness, we generated multiple SNP datasets optimized for specific objectives. Within each dataset, individuals with excessive missing data were removed using the –mind filter based on the average missingness of the dataset (--mind 0.3-0.5).

After filtering in PLINK, we transformed the datasets into genetix files for further analysis using PGDSpider (v 2.1.1.0). We further filtered SNP datasets for deviations from Hardy Weinberg Equilibrium via the “out-all” method (removes loci that deviate from HWE in all populations) using the R package *DartR* (Gruber et al. 2018).

### 2.5. Texas vs. North Carolina

To identify broad-scale genetic differentiation between Texas (TX) and North Carolina (NC), we analyzed all individuals from both states grouped by sampling year (e.g. NC 2014, TX 2015). We then partitioned by year (i.e. TX vs. NC 2014, TX vs. NC 2015, TX vs. NC 2016).

We calculated pairwise measures of genetic differentiation (*F_ST_*) using the R package *hierfstat* (v. 05-11; Goudet 2005) for each sampling year (2014-2016) using the command “pairwise.WCfst” (Cockerham and Weir 1984). We then used “boot.ppfst” to calculate *F*_ST_ confidence intervals (CI=0.025, 0.975) using 1000 bootstraps. The “boot.ppfst” function repeatedly resamples loci to generate empirical distributions of pairwise *F*_ST_ estimates.

### 2.6. Texas

The dataset that included all Texas locations and years consisted of 38,013 loci with an overall genotyping rate of 0.90. We then analyzed each sample year separately (2014, 2015, 2016). For each year, we calculated genetic summary characteristics, including observed heterozygosity (*H_o_*), expected heterozygosity (*H_E_*) and the inbreeding coefficient (*F_IS_*), using the *basic.stats* function in *hierfstat* (Goudet 2005).

To identify genetic clusters and estimate cluster membership probabilities within Texas, we performed principal components analysis (PCA) for each sampling year in R using the packages *adegenet* (v. 1.3-1; Pritchard et al. 2000) and *Ade4* (v. 1.7-23; Dray & Dufour 2007). We used PCA analyses to determine clustering as it does not require *a priori* group assignment. We retained the first two principal components of each PCA because they explained the greatest proportion of genetic variance and no additional structure was apparent in subsequent components. We then produced scatter plots using *ggplot2* in R (Wickham 2016).

For the one location in Texas, Bastrop Bayou, that we had yearly sampling (2014, 2015, and 2016), we calculated pairwise *F_ST_* values in *hierfstat* and conducted a PCA in *adegenet* to assess temporal differentiation within this location.

### 2.7. North Carolina

To investigate population structure among sampling locations within NC, individuals were grouped by location and year. We ran the *populations* pipeline in STACKS with a stricter locus sharing criterion (-p 2) to remove loci present in only a single population and reduce the influence of rare or location-specific variants. After filtering, the NC dataset consisted of 25,085 loci.

To produce genetic summary statistics, we grouped the dataset by each region (Albemarle, Pamlico Sound, Neuse River, New River) and year, and calculated statistics using *basic.stats* in *hierfstat*.

Then, to determine population genetic structure among specific locations in NC, we grouped the dataset by specific sampling location and year and used this dataset to calculate pairwise *F_ST_* values. Although some sampling locations within NC contained a small number of individuals (n= 4-5), this limitation is mitigated by the large number of SNPs analyzed (25,085). Previous studies have shown that increasing the number of loci improves the accuracy of population genetic inference and can compensate for a small number of individuals (e.g. Willing et al. 2012; Nazareno et al 2017; McLaughlin & Winker 2020). To confirm patterns of genetic structure, we also calculated pairwise *F_ST_* among combined regions and years (Albemarle, Pamlico Sound, Neuse River, New River), which increased sample sizes by pooling individuals from multiple locations within each region (Table S4). We also conducted multivariate and clustering analyses including principal components analysis (PCA) and ADMIXTURE (v. 1.3; Alexander et al. 2009) for specific NC sampling locations and years. To determine the optimal value for K, we generated standard error scores using cross-validation for K=1-n (where n is the total number of *a priori* sampling locations in the dataset) and selected the K value with the lowest cross-validation error. We generated plots in R using *ggplot2* from ADMIXTURE Q matrices with individuals ordered North to South by location.

To assess temporal genetic structure within regions, we analyzed locations sampled in at least three years. These locations included: one location in the Pamlico Sound Region - Swanquarter (2014, 2015, 2016); three locations in the Neuse River Region - Hancock Creek (2014, 2016, 2022, 2023), Slocum Creek (2014, 2016, 2023), and Clubfoot Creek (2014, 2016, 2022, 2023); and two locations in the New River Region - Mill Creek (2014, 2015, 2016) and Virginia Creek (2014, 2016, 2023). Note that we included the two sites sampled in 2022 in the temporal datasets for Hancock and Clubfoot Creeks in the Neuse River Region. We calculated pairwise *F_ST_* for each temporal dataset in *hierfstat* and produced a PCA in *adegenet*.

## 3. Results

### 3.1 Broad geographic-scale differentiation - North Carolina and Texas

We found consistent, and significant genetic differentiation between the two ocean basins in Texas and North Carolina in all three sampling years (*F_ST_* values = 0.0467 (2014), 0.0542 (2015) and 0.0620 (2016); Table S2). All values were significant based on confidence intervals produced with 1000 bootstraps.

### 3.2 Texas

#### 3.2.1. Genetic characteristics

Genetic diversity in Texas was similar across the limited sampling years (2014–2016). We found expected heterozygosity (*H_E_*) ranged from 0.1224 to 0.1362 and all years showed positive inbreeding coefficients (*F_IS_* = 0.0949 to 0.1060; Table 1). This indicated a slight deficit of heterozygotes relative to Hardy–Weinberg expectations.

**Table 1.** Genetic statistics for Texas samples by year. N represents the number of individuals sampled, Ho is observed heterozygosity, He is expected heterozygosity, and Fis is the inbreeding coefficient calculated from genome-wide SNP data for 2014–2016.

|  | <u>N</u> | <u>H<sub>O</sub></u> | <u>H<sub>E</sub></u> | <u>F<sub>IS</sub></u> |
| --- | --- | --- | --- | --- |
| **Texas 2014** | 34 | 0.1218 | 0.1362 | 0.1060 |
| **Texas 2015** | 24 | 0.1108 | 0.1224 | 0.0952 |
| **Texas 2016** | 9 | 0.1185 | 0.1309 | 0.0949 |

#### 3.2.2. Spatial structure in Texas

The first two principal components of the Texas PCA explained 2.6% and 2.1% of the total genetic variation (Fig. 3A). The PCA revealed limited genetic differentiation among most Texas sampling locations and years. Individuals from East Bay (2014), Chocolate Bayou (2014), and Bastrop Bayou (2014–2016) largely overlapped near the center of the plot, indicating little genetic structuring among these sites or across Bastrop Bayou sampling years. In contrast, individuals from the 2016 hatchery population formed a distinct cluster separated from the wild samples along PC1.

**Fig 3.**
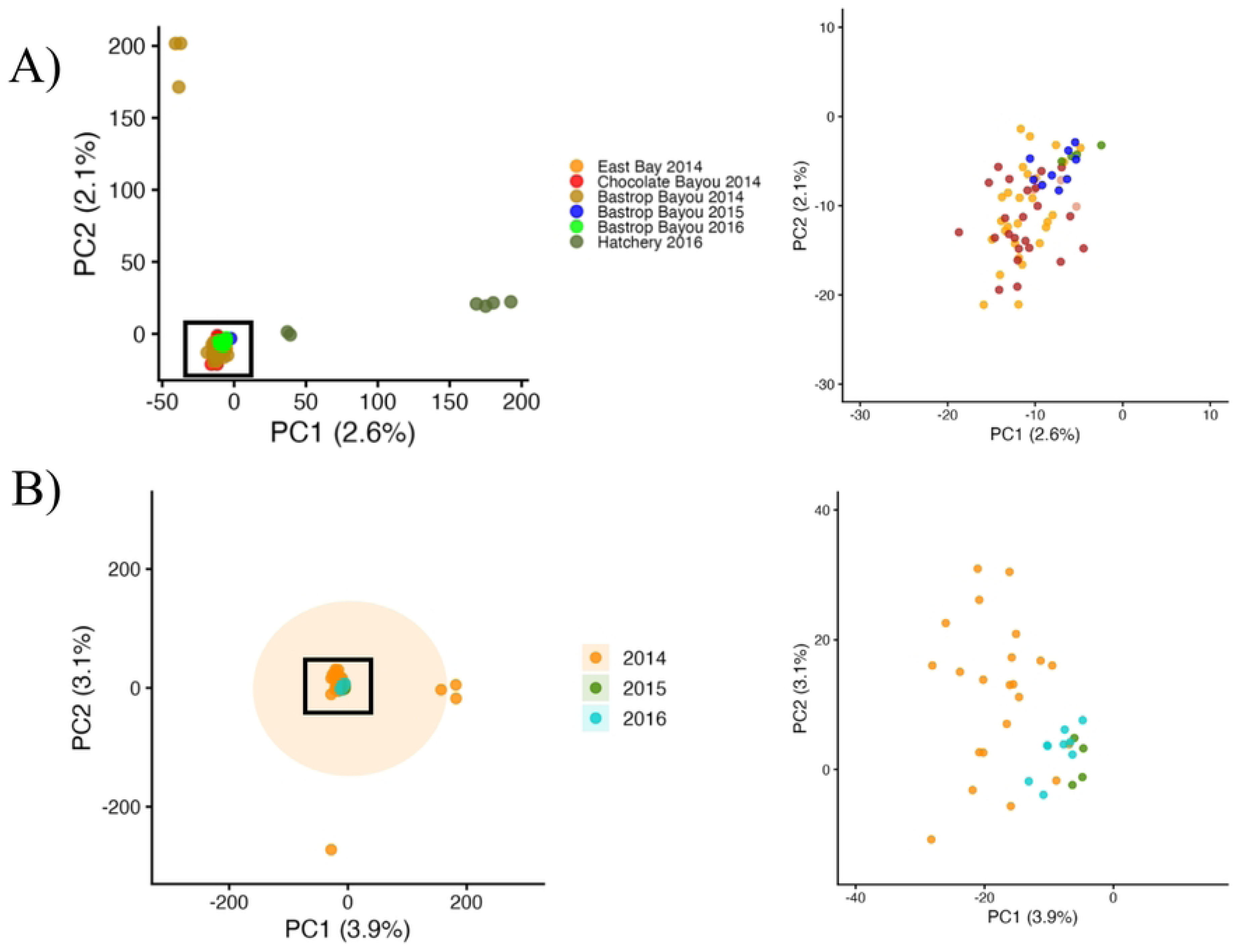
Principal component analysis (PCA) of southern flounder (*Paralichthys lethostigma*) from Texas. (A) PCA including all sampled locations and years: East Bay (2014), Chocolate Bayou (2014), Bastrop Bayou (2014–2016), and hatchery (2016). (B) PCA including only Bastrop Bayou samples from 2014–2016. Left panels show the full PCA, and right panels show a magnified view of the central cluster (boxed). Points represent individuals and are colored by location (A) or year (B). Axes indicate the percent variance explained by each principal component.

#### 3.2.3. Temporal structure in Bastrop Bayou

The temporal PCA of Bastrop Bayou, TX, showed that 2015 and 2016 clustered tightly together, while 2014 showed a high degree of individual variation along both the PC1 and PC2 axes (3.9% and 3.1%; Fig. 3B). In contrast, we found significant differentiation between 2015 and 2016 based on pairwise *F_ST_* analysis. *F_ST_* values ranged from 0.0002 - 0.0035, and the comparison between 2015 and 2016 was significant based on 95% confidence intervals (Table S7G).

### 3.3 North Carolina

#### 3.3.1 Regional Genetic Characteristics

Genetic diversity was similar across regions and years (Table 2). Expected heterozygosity (*H_E_*) ranged from 0.1413 to 0.1614, with the highest value observed in Neuse River (2016). Across all populations, *H_E_* was slightly higher than *H_O_*, resulting in consistently positive inbreeding coefficients (*F_IS_* = 0.070 to 0.105).

**Table 2.** Genetic statistics for North Carolina samples by region and year. N represents the number of individuals sampled, Ho is observed heterozygosity, He is expected heterozygosity, and Fis is the inbreeding coefficient calculated from genome-wide SNP data for 2014–2016 and 2023.

|  | <u>N</u> | <u>H<sub>O</sub></u> | <u>H<sub>E</sub></u> | <u>F<sub>IS</sub></u> |
| --- | --- | --- | --- | --- |
| **Albemarle 2023** | 61 | 0.1407 | 0.1565 | 0.1013 |
| **Pamlico 2014** | 21 | 0.1480 | 0.1603 | 0.0763 |
| **Pamlico 2015** | 22 | 0.1330 | 0.1454 | 0.0856 |
| **Pamlico 2016** | 9 | 0.1485 | 0.1604 | 0.0745 |
| **Neuse 2014** | 20 | 0.1474 | 0.1585 | 0.0701 |
| **Neuse 2015** | 26 | 0.1294 | 0.1413 | 0.0842 |
| **Neuse 2016** | 29 | 0.1489 | 0.1614 | 0.0778 |
| **Neuse 2023** | 12 | 0.1462 | 0.1581 | 0.0757 |
| **New River 2014** | 18 | 0.1488 | 0.1605 | 0.0728 |
| **New River 2015** | 7 | 0.1519 | 0.1519 | 0.0799 |
| **New River 2016** | 26 | 0.1479 | 0.1600 | 0.0752 |
| **New River 2023** | 15 | 0.1544 | 0.1544 | 0.1053 |

#### 3.3.2 Overall Spatial Structure in North Carolina by Year

Within North Carolina, we found significant genetic differentiation that varied among sampling locations and years. Full pairwise *F*_ST_ matrices are found in supplementary tables S3-S6. Although the observed pairwise *F*_ST_ values were small in magnitude, these values were expected given the marker type in this study and supporting consistent but spatially variable genetic structure in this region. Because our sampling sizes were small in some locations, we assessed pairwise structure among broad regions and years and found that almost all pairwise comparisons were significant (Table S8).

To better understand these patterns, we examined the genetic structure within each year. In 2014, we did not find significant pairwise *F*_ST_ values (Table S3), and the PCA revealed minimal genetic clustering among sampling locations (Fig. 4A). The first two components of the PCA explained 2.1% and 2.0% of the total genetic variance, respectively. Individuals from most populations showed substantial overlap in multivariate space, indicating limited genetic differentiation. In the New River Region, samples from Virginia Creek showed greater dispersion along PC1 and PC2 and included several highly divergent individuals relative to other populations.

**Fig 4.**
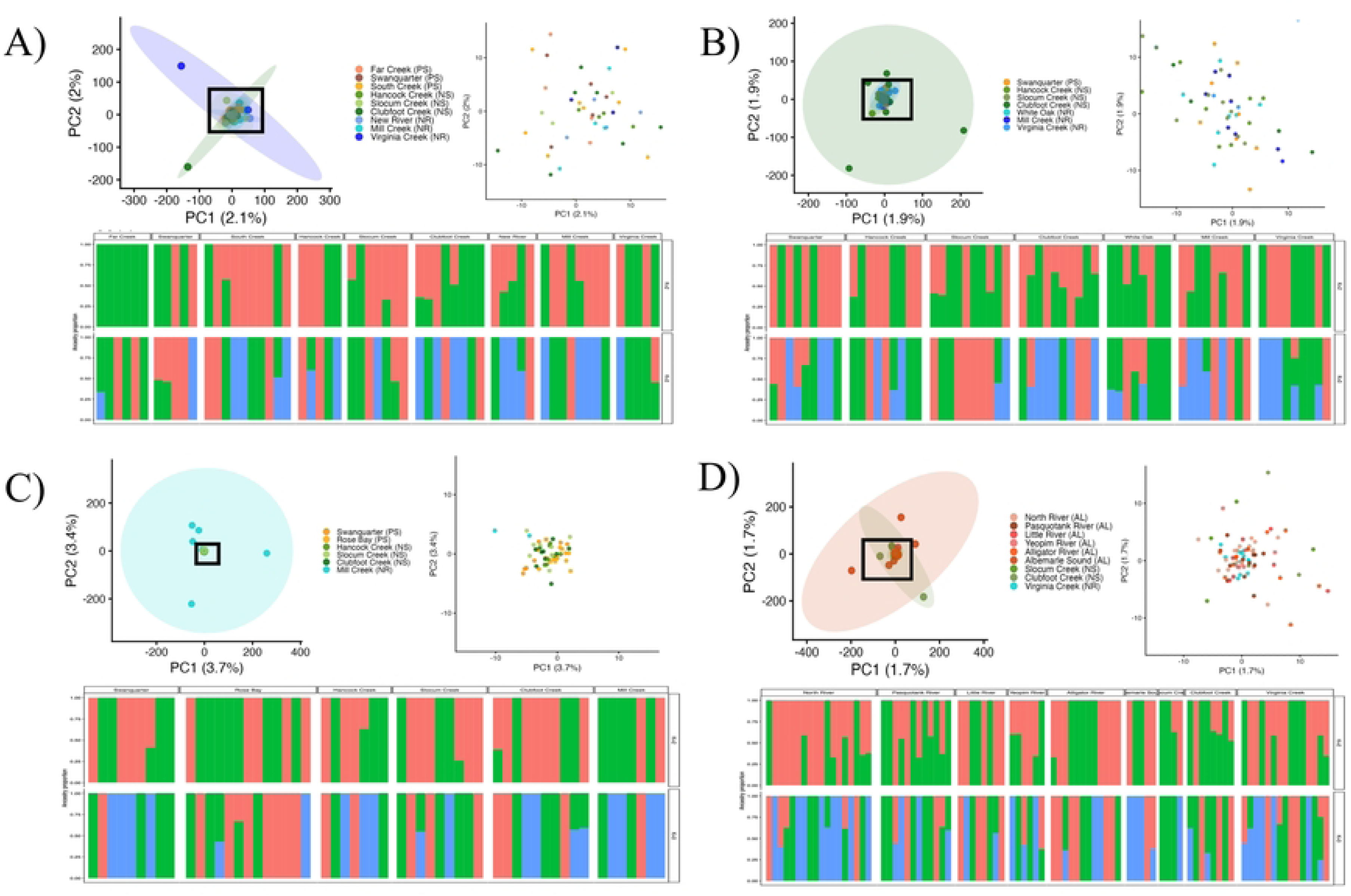
Population structure of southern flounder (*Paralichthys lethostigma*) sampled from North Carolina estuaries across four years. (A) 2014, (B) 2015, (C) 2016, and (D) 2023. For each year, the upper panels show a principal component analysis (PCA) of individuals by sampling location, with the full PCA on the left and a magnified view of the central cluster (boxed) on the right. Axes indicate the percent variance explained by each principal component. The lower panels show admixture results for K=2(top) and K=3(bottom), with each vertical bar representing an individual and colors indicating inferred ancestry proportions.

In 2015, we found significant pairwise *F*_ST_ values at five of the 15 comparisons (Table S4), which was supported by the results of our PCA. Of the five significant comparisons in the pairwise *F*_ST_ analysis, four included the Mill Creek site in the southernmost New River Region. We also found significant pairwise *F_ST_* between two sites in Pamlico Sound: Rose Bay and Swanquarter. The results of the PCA in 2015 explained 3.7% and 3.4% of variation, respectively (Fig 4B). We found notable overlap of all sampling sites except samples from Mill Creek, which were highly dispersed across the ordination space and included several individuals strongly diverged from all other populations in the PCA.

In 2016, we did not find significant pairwise *F_ST_* values (Table S5), and the results of the PCA supported a lack of population genetic structure as most sampling sites clustered tightly (Fig. 4C). However, the Clubfoot Creek location did include some divergent individuals that dispersed along PC1 and PC2. Both axes explained 1.9% of the total genetic variance.

In 2023, we included samples from Albemarle Sound Region, which was the northernmost region sampled. With the addition of this northernmost region, we found greater genetic differentiation among sites. We found significant pairwise *F*_ST_ values at 13 out of 36 comparisons (Table S6). Of these 13 comparisons, six were between Albemarle and Neuse locations and seven were among Albemarle sampling locations. The results of the PCA showed both axes explained 1.7% of the total variance (Fig. 4D). While most individuals from all sampling locations clustered tightly, samples from Albemarle Sound and Clubfoot Creek (Neuse) showed some individual divergence along the PC1 and PC2 axes.

The results of the ADMIXTURE analysis were similar among all years (Fig 4A-D). Across all years, cross-validation (CV) error was lowest at K=1 and increased at higher K values, consistent with panmixia and a lack of genetic structure across *a priori* sampling locations. Although cross-validation supported K=1 in all years, admixture plots showed some individual-level variation in ancestry proportions.

#### 3.3.3 Temporal Variation of Specific NC Sampling Sites

When we analyzed year to year variation within a site, we found significant genetic differences across years within sites in our pairwise *F_ST_*analysis that was consistent with the results of our PCAs.

*Pamlico Region*. For Swanquarter, we found significant genetic differentiation between years in our pairwise *F_ST_* analysis at two of the three comparisons: 2015 compared to either 2014 (*F_ST_*= 0.0056) or 2016 (*F_ST_* = 0.0027; Table S9A). For the PCA, we found PC1 and PC2 explained 5.5% and 5.6% of the variation. Samples from 2014 and 2015 formed a tight cluster, while 2016 samples showed slightly greater dispersion along both PC1 and PC2 (Fig. 5A).

**Fig 5.**
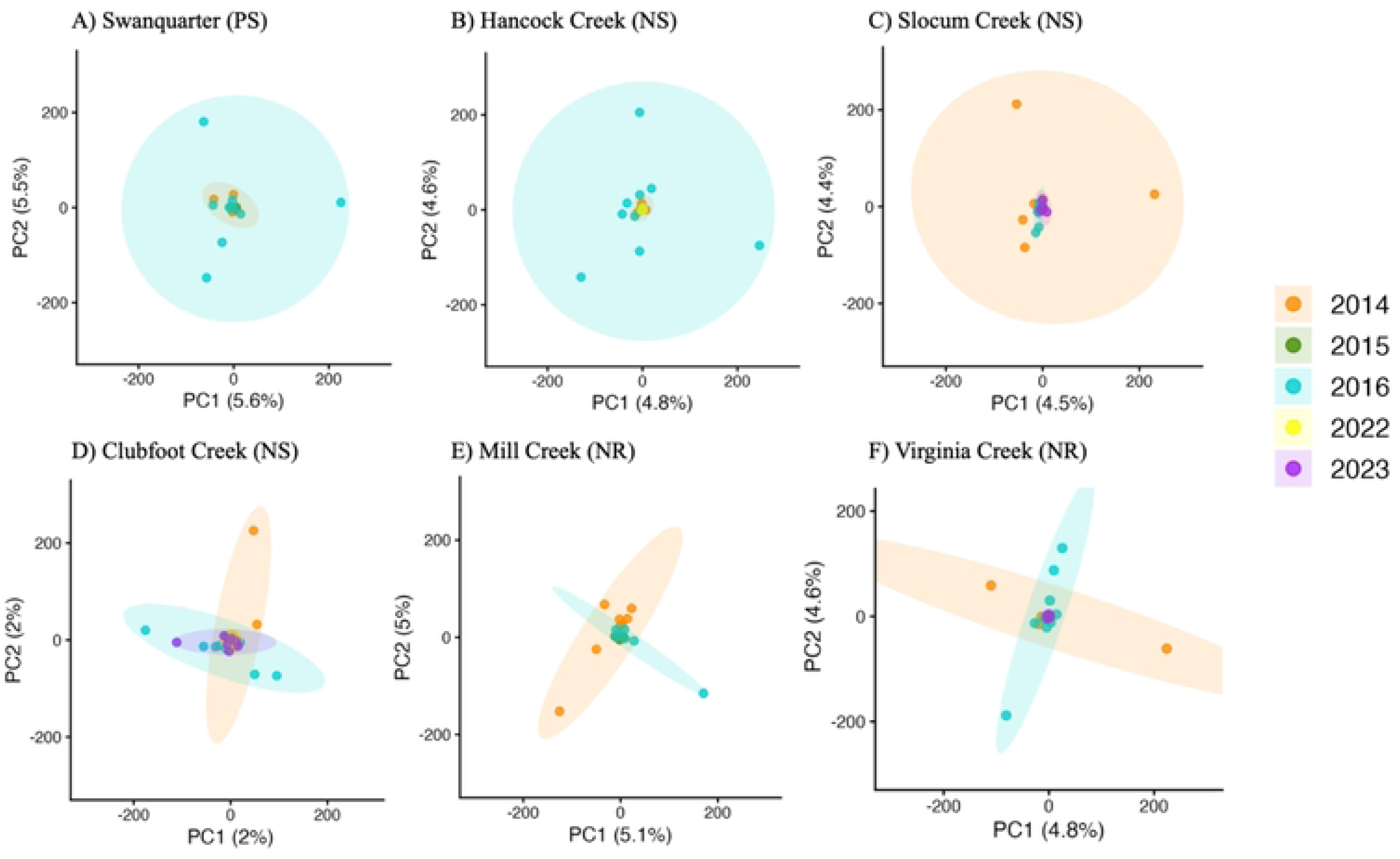
Principal component analyses (PCA) of southern flounder (*Paralichthys lethostigma*) used to examine temporal population structure within North Carolina estuaries. Sampling locations are arranged geographically from north to south (top left to bottom right): (A) Swanquarter (Pamlico Sound; PS), (B) Hancock Creek (Neuse River; NS), (C) Slocum Creek (Neuse River; NS), (D) Clubfoot Creek (Neuse River; NS), (E) Mill Creek (New River; NR), and (F) Virginia Creek (New River; NR). Points represent individuals colored by sampling year (2014, 2015, 2016, 2022, and 2023), with ellipses indicating dispersion within years. Axes show the percent variance explained by the first two principal components.

*Neuse River Region.* In this region we had three sites that had multiple years sampled. We found significant genetic differentiation among years at Hancock Creek at four of six pairwise comparisons (*F_ST_* = 0.0014 - 0.0040; Table S9B). The PCA (PC1: 4.8%, PC2: 4.6%) showed overlap between 2014, 2015 and 2022, while 2016 individuals clustered separately showing high divergence. We also found significant genetic differentiation among years in Clubfoot Creek at eight of ten pairwise comparisons (*F_ST_* = 0.0016 - 0.0036; Table S9C). We found greater divergence among individuals at Clubfoot Creek compared to Hancock Creek in the PCA analysis (Fig. 5). The Clubfoot Creek PCA (PC1: 5.6%, PC2: 5.5%) showed that, while most individuals cluster tightly, 2014, 2015, and 2016 all contain highly differentiated individuals, with 2016 showing the highest degree of divergence. At the third location, Slocum Creek, we found significant genetic differentiation among years at five of the six pairwise *F_ST_* comparisons (*F_ST_* = 0.0027 - 0.0035; Table S9C), and the PCA (PC1: 4.5%, PC2: 4.4%) showed 2015, 2016 and 2023 clustered tightly with some separation between 2016 and 2023. In addition, 2014 contained highly divergent individuals on both axes.

*New River Region*. In this southernmost region, we found greater genetic structure among years at the two sites we had yearly sampling than the other two regions. At Mill Creek we found significant genetic differentiation among years in two of three pairwise *F_ST_* comparisons (0.0031-0.0039; Table S9E). The PCA showed that 2023 clustered together tightly, while 2014 and 2016 contained highly divergent individuals that clustered on separate axes (Fig. 5E). In Virginia Creek, no pairwise comparisons were significant (Table S9F). The PCA showed a similar pattern to Mill Creek, with 2015 clustered tightly, while 2014 and 2016 showed higher divergence on separate axes.

## 4. Discussion

Our study evaluated spatial and temporal genetic structure of juvenile southern flounder from two different basins in its range, specifically in Texas and North Carolina, with a particular emphasis on the poorly studied North Carolina region in the Atlantic where there has been a substantial decline in flounder populations (Flowers et al. 2019). Previous studies on the southern flounder have either pooled samples across years or lacked fine-scale spatial sampling at scales likely corresponding to the contributions from individual spawning stocks. By incorporating both broad and fine spatial scales and sampling across multiple years, we aimed to capture recruitment patterns associated with discrete yearly cohorts of settled juveniles in estuarine locations rather than signals averaged across multiple cohorts. This approach enabled us to resolve spatial and temporal variation in spawning stock contributions among estuaries and years.

At a broad scale, our analyses revealed clear genetic differentiation between Texas and North Carolina, consistent with previous studies documenting large basin genetic structure between Gulf of Mexico and Atlantic populations (Anderson et al. 2012; Wang et al. 2015; O’leary et al. 2021). Within Texas and North Carolina, we detected fine-scale genetic structure that varied in strength across sampling years, suggesting temporal shifts in the genetic composition of recruiting juveniles from the adult spawning stock. In North Carolina, we observed year-to-year variation in genetic composition among individuals within sampling sites, indicating that local nurseries may receive recruits from multiple spawning cohorts. Together, our findings highlight the importance of including both spatial and temporal sampling when evaluating population structure to more effectively resolve spawning stock contributions to estuarine recruitment.

### 4.1 Broad-scale differentiation between North Carolina and Texas

The significant genetic differentiation between southern flounder from different basins, The Atlantic (North Carolina) and Gulf of Mexico (Texas), supports earlier studies using otolith chemistry, microsatellites, AFLPs, and SNPs, all of which suggest the Gulf and Atlantic represent distinct stocks (Anderson et al. 2012; Anderson & Karel 2012; Wang et al. 2015; O’Leary et al. 2018). Anderson et al. (2012) proposed that this divergence reflects independent post-Last Glacial Maximum expansion of southern flounder populations in the Gulf of Mexico and Atlantic, resulting in long-term isolation between basins. This Atlantic-Gulf split is a common pattern among coastal marine species, including red drum, an estuarine-dependent species with comparable life history traits (Hollenbeck et al. 2018). Currently, Atlantic and Gulf flounder are managed as separate stocks, and our findings provide additional support for maintaining this management framework.

### 4.2 Regional differentiation in Texas

Within Texas, we did not find significant genetic divergence among sites within a year; however, samples from Bastrop Bayou exhibited significant interannual genetic differentiation suggesting that larval recruitment or spawning stock composition at this location varies among years. This variability likely reflects the dynamic oceanographic processes in the Gulf of Mexico. Fluctuations in wind-driven circulation, freshwater influx from river discharge, and mesoscale features such as eddies can influence larval transport, retention, and connectivity among estuaries (Taylor et al. 2010). Combined with variability in spawning time and reproductive success, these processes can result in shifting contributions from spawning stock among years, resulting in variable temporal genetic structure. Our findings provide a temporal context to what was reported in O’Leary et al. 2021, who found genetic heterogeneity within the Gulf of Mexico. While our Gulf sampling was limited to sites within Texas, we found temporal differences among years that support the existence of fine-scale, dynamic population structure within the Gulf of Mexico.

Our analysis revealed that individuals from the hatchery were genetically distinct from wild-caught juveniles, suggesting that the hatchery broodstock may not fully represent the local spawning stock. This is likely due to founder effects, long-term inbreeding of hatchery individuals, and/or non-local sourcing. However, the genetic diversity (*H_E_*) was not significantly reduced relative to wild populations; therefore, this pattern likely reflects a distinct lineage rather than a recent bottleneck. While the introduction of genetically distinct individuals could, in some cases, increase genetic diversity in wild populations, such outcomes depend on the composition of the hatchery stock and the degree of selection for hatchery conditions. Hatchery lines affected by founder effects, inbreeding, or selection may be maladapted to natural environments, and their introduction can reduce the overall fitness of the population they are introduced to. Similar challenges have been documented in other fish species such as salmonids (Araki et al. 2007; Christie et al. 2012), where hatchery releases led to reduced fitness or altered population structure in wild populations. These results highlight the importance of carefully evaluating broodstock sourcing and management in stock enhancement programs to ensure that efforts support the genetic integrity and adaptive potential of wild southern flounder populations.

Overall, our results suggest a pattern of high connectivity within Texas accompanied by localized and temporally variable genetic structure, likely driven by recruitment processes rather than persistent barriers to gene flow. While most studies have found limited genetic differentiation in the Gulf of Mexico (Anderson & Karel, 2012), some evidence supports the presence of finer-scale structure (Wang et al. 2015). Both studies included a mixture of life stages (i.e. juveniles and adults or multiple age classes) which may integrate signals across cohorts and years and therefore reflect a time-averaged view of connectivity. In contrast, our focus on juveniles likely enhances the detection of cohort-specific genetic structure. Although our sampling was limited to Texas and did not reveal consistent large-scale structure, dynamic patterns may occur elsewhere in the Gulf. Expanded spatial and temporal sampling will be necessary to more fully resolve southern flounder population dynamics across this region.

### 4.3 Regional differentiation in North Carolina

Within North Carolina, we found significant fine-scale genetic structure among estuarine sites that varied among years. Some individual sampling locations showed greater genetic divergence across years than other sampling locations within the same year, indicating temporal variability in population composition. This inconsistent temporal structure likely reflects interannual variation in the amount and/or source of larval recruitment. During the pelagic larval stage, survival and settlement are highly sensitive to environmental variability including wind, currents and freshwater discharge (Hare & Cowen 1996; Pineda et al. 2007; Taylor et al. 2010). Because larval mortality is high, even minor shifts in these conditions can lead to sweepstakes reproductive success, in which a limited number of adults contribute disproportionately to the reproductive gene pool (Hedgecock 1994). This process, also described as chaotic genetic patchiness (CGP; Hedgecock 1994), can generate spatial and temporal genetic differences among juveniles even when long-term population structure is weak or absent. Similar patterns have been documented in numerous pelagic-larval species, including the bicolor damselfish (Christie et al., 2010) and the Antarctic limpet (Vendrami et al. 2021), where temporally unstable genetic structure has been observed.

In addition to this temporal variability, we also observed substantial genetic variation among individuals within the same sampling locations. This suggests that estuarine nursery habitats may receive recruits from multiple spawning groups or source areas. Such mixed recruitment implies that individuals within a single location may originate from different parental populations or spawning events, further contributing to the subtle and temporally variable genetic patterns observed among sites. These findings highlight the importance of incorporating multi-year genetic data when evaluating population structure, as conclusions drawn from a single sampling year may represent only a snapshot of a dynamic system, and could lead to uniformed management or broodstock sourcing decisions. It also suggests that management should focus on maintaining large and genetically diverse spawning stocks to maintain viable population growth.

Previous studies found little evidence of population genetic structure within the Atlantic (Anderson et al. 2012; Wang et al. 2015; O’Leary et al. 2021). However, these studies were limited by lower marker resolution, reduced fine-scale sampling, and a lack of temporal replication. Earlier work using microsatellites or allozymes, as well as SNP based studies with fewer loci (O’Leary et al., 2018), may have lacked the power to detect weak or transient genetic structure. Additionally, sampling across multiple years likely obscured temporal variation by averaging signals across cohorts. In contrast, our use of a larger genome-wide SNP dataset (∼25,000 SNPs), combined with sampling at spatial scales relevant to individual estuaries and across discrete years, increased our ability to detect subtle patterns of differentiation. The inconsistent fine-scale structure we observed within North Carolina likely reflects temporally variable recruitment and shifting spawning stock contributions rather than stable subdivision. As such, genetic structure at this scale is likely ephemeral and cohort-dependent, making it difficult to detect in studies that average across years or sample on broader spatial scales. These results suggest that the apparent lack of structure reported in earlier studies may reflect limitations in sampling design and marker resolution rather than true panmixia within Atlantic populations.

Within North Carolina, we detected several notable regional patterns. The southern New River region exhibited more consistent differentiation from other locations, potentially reflecting environmental influences on larval dispersal, recruitment, or local selection. Honeycutt et al. (2019) found higher temperatures in the New River region were associated with more male-biased sex ratios in southern flounder, likely due to temperature-dependent sex determination (TSD). This suggests that temperature-driven selection could be driving genetic differentiation in this region. Asynchronous reproduction may also be driving these differences. Honeycutt et al. (2019) also reported that some New River region fish had similar growth patterns to fish sampled from South Carolina, where migration and spawning occur earlier (Wenner 1990), while others were smaller, suggesting a later spawning event. These observations indicate that some New River fish may be moving offshore to spawn earlier in the season than northern regions, leading to temporal spawning stock structure and genetic differentiation. Additional sampling from the New River region is needed to better understand how environmental conditions and reproductive timing shape genetic structure in this region. As ocean temperatures continue to rise across the southeastern Atlantic (Cheng et al. 2023), further research with expanded sampling in this region will be important to determine how temperature-driven processes influence selection and reproductive behavior in southern flounder populations at local levels.

Albermarle Sound, the northernmost estuary in this study that we sampled in 2023, exhibited strong differentiation among sites both within and outside the Albemarle region. With only a single inlet connecting Albemarle to the broader Atlantic via Pamlico Sound, opportunities for gene flow may be constrained, leading to local differentiation even across small spatial scales within this Sound (Cowen et al. 2000). These findings raise the possibility that restricted oceanographic connectivity may contribute to the pronounced genetic structure observed in this region. To determine the mechanism causing increased genetic differentiation, and whether this is a consistent pattern in Albemarle Sound, will require additional sampling, including multi-year sampling.

### 4.4. Management and Conclusions

From a management perspective, maintaining a large, diverse spawning stock is critical, as it influences both the stability and diversity of recruitment (Anderson et al. 2008). Larger spawning stock populations are more likely to include individuals that spawn across a range of locations and times, increasing the diversity of reproductive contributions to each cohort (Schindler et al. 2010). This diversity is particularly important in dynamic coastal systems, where environmental variability, such as changes in temperature, salinity, and circulation can differentially affect larval survival and dispersal over short spatial and temporal scales (Cowen & Sponaugle 2009). By supporting a wide range of spawning times and locations, large spawning stocks enhance the potential for successful recruitment under varying conditions, thereby increasing population resilience to environmental change. Incorporating genetic data across multiple years will improve the ability to detect these dynamics, avoid bias toward single cohorts or recruitment events, and support management strategies that sustain both connectivity and adaptive potential (Benestan 2019).

Overall, our results demonstrate that southern flounder population structure is shaped by dynamic interactions among spatial, temporal, and environmental processes, leading to patterns that are often subtle and cohort dependent. Recognizing this variability is critical for accurately interpreting connectivity and recruitment across the species’ range. Our findings highlight the value of integrating fine-scale, multi-year genetic data to capture these dynamics and avoid misleading conclusions based on single-year or coarse-scale sampling. The use of high-resolution genomic approaches, such as ddRAD sequencing, enables the analysis of thousands of SNP markers and further enhances the ability to detect variation in population structure. Moving forward, expanded spatial and temporal sampling, coupled with environmental data, will be essential for linking oceanographic processes, reproductive behavior, and genetic structure.

These efforts will strengthen management strategies that maintain population connectivity, preserve adaptive potential, and support the long-term resilience of this commercially important finfish species.

## 7. Author Contributions

**S.P. Harned:** Conceptualization, Data Collection, Data Curation, Methodology, Formal Analysis, Writing - Original Draft, Writing - Editing & Reviewing. **J.L. Mankiewicz:** Data Collection, Writing - Editing & Reviewing. **R. J. Borski:** Formal Analysis, Writing - Editing & Reviewing. **J. Godwin:** Formal Analysis, Writing - Editing & Reviewing. **M.O. Burford Reiskind:** Conceptualization, Methodology, Formal Analysis, Resources, Writing - Editing & Reviewing.

## 8. Data Accessibility

Raw sequence data have been deposited in the NCBI Sequence Read Archive under BioProject accession PRJNA1295924. BioSample accessions for each individual are listed in the associated metadata table. Ped and .map files for each dataset, containing individual SNP genotypes prior to PLINK filtering, are available on Dryad. Gtx files used for downstream analysis post-filtering are also available on Dryad (Dataset DOI: 10.5061/dryad.sn02v6xhq). Data will be released upon publication.

## 10. Acknowledgements

We would like to thank the NC Division of Marine Fisheries, specifically T. Moore, D. Zapf, T. Wadsworth, C. Stewart, M. Hamric, A. Markwith, and H. White, and Texas Parks and Wildlife, specifically Lindsay Campbell, for providing us with flounder samples. We would also like to thank Caitlin Mothes and Emma Wallace for completing the DNA extractions and ddRADseq library building for the 2014, 2015, and 2016 years.

